# Semi-quantitative Classification of HIV-1 Nucleic Acids Using ResNet Image Analysis of Discretized Isothermal Amplification Reactions in a Microfluidic Chip

**DOI:** 10.64898/2026.06.24.734232

**Authors:** CD Martin, NC Benson, NS Gummalla, KN Shimazu, AT Bender, DAC Beck, JD Posner

**Affiliations:** Department of Chemical Engineering, University of Washington, Seattle, Washington, USA; eScience Institute, University of Washington, Seattle, Washington, USA; Department of Mechanical Engineering, University of Washington, Seattle, Washington, USA; Department of Family Medicine, University of Washington, Seattle, Washington, USA

## Abstract

Isothermal nucleic acid amplification tests enable rapid and decentralized molecular diagnostics but often lack robust quantitative readouts compared to quantitative PCR. Here, we present a semi-quantitative nucleic acid measurement approach using machine learning to extract spatiotemporal features from real-time fluorescence imaging of rapid isothermal amplification reactions in microfluidic chips. A convolutional neural network was trained on multiple images sampled throughout a chip-based recombinase polymerase amplification reaction to classify samples into clinically relevant or logarithmically spaced concentration ranges spanning five orders of magnitude. The clinical classification model achieved 94.6% accuracy, and the logarithmic model achieved 92.7% accuracy, with most errors occurring between adjacent concentration categories. By learning spatiotemporal patterns of fluorescence development rather than relying on explicit feature extraction, the model remained accurate at both high and low nucleic acid concentration regimes where other quantitative isothermal molecular tests struggle. This approach enables automated interpretation of amplification reactions and extends the usable dynamic range of the assay. These results demonstrate that integrating machine learning with image-based amplification methods can support rapid semi-quantitative molecular testing and may facilitate broader deployment of nucleic acid diagnostics outside centralized laboratory settings.

**Author summary:** Many rapid nucleic acid testing methods for infectious diseases are simple to run but struggle to measure how much genetic material is present, which limits their usefulness in clinical decision-making. In our work, we study a technique that produces visible fluorescent patterns during nucleic acid amplification reactions. Traditionally, the amount of nucleic acids present are measured by counting individual bright spots, but this becomes difficult when the target nucleic acid concentration is high and the spots merge together.

We developed a machine learning approach that models how the fluorescence pattern changes over time. By analyzing a sequence of images from each reaction, our model can assign samples to concentration ranges across a wide span. This allows us to extract meaningful information even when traditional analysis methods break down. Because this approach works with simple imaging systems and does not require complex equipment, it could help support more informative and accessible diagnostic testing in point-of-care and low-resource settings.

## Introduction

The clinical diagnostics field is undergoing a shift away from centralized laboratory tests that can take hours or days to point-of-care (POC) tests that are more accessible, affordable, and deliver accurate results within minutes. Artificial intelligence (AI) and machine learning (ML) are accelerating this transition by improving the performance and capabilities of existing diagnostic technologies, such as rapid lateral flow assays and nucleic acid amplification tests (NAATs) [1–6]. Some POC tests leverage smartphones or compact readers to image, analyze, and display results to end users. Recent studies have shown that on-device ML algorithms can be integrated with NAATs to automate result interpretation on smartphones and compact readers [7–10]. For example, ML has been shown to result in more accurate determinations of results in cases of low signal output or where human or threshold-based reads are error prone by analyzing real-time fluorescence trajectories with RNN/LSTM/Transformer models [4,11,12]. These advances highlight the potential of ML-enhanced POC NAATs to improve analytical sensitivity, shorten runtime, enable multiple target detection, and support semi-quantitative nucleic acid measurement.

The NAAT field has rapidly advanced beyond centralized, gold standard, Polymerase Chain Reaction (PCR) [5,6]. Isothermal methods, such as Recombinase Polymerase Amplification (RPA) and Loop Mediated Amplification (LAMP), have enabled decentralized nucleic acid testing by reducing hardware burdens, shortening runtimes, offering flexible readout modalities, and achieving high sensitivity and specificity [13–15]. A notable drawback of these methods is that they are not inherently quantitative like PCR, where amplification events are synchronized with each thermal cycle. Studies have explored quantitative, isothermal NAATs using bulk fluorescence or time to a predefined fluorescence threshold, but these approaches have limited reproducibility and dynamic ranges [11,13–19]. Isothermal digital NAATs have demonstrated absolute nucleic acid quantification in specialized microchips where reactions are partitioned into droplets or microwell arrays and binary amplification results of individual wells are correlated directly with input nucleic acid concentration [20–24]. ML and AI tools, such as Meta’s zero-shot Segment Anything Model, have been applied for analyzing digital NAAT data sets where it is critical to accurately count the number of positive amplification events across an array of hundreds or thousands of droplets or microwells [25–27]. In moving towards decentralized quantitative testing, isothermal digital LAMP assays have leveraged deep learning for droplet segmentation and identification, color coding, and precipitate characterization for quantitative assays that offer multiplexing functionality [28–31].

We previously introduced a digital RPA technique, called Amplification Nucleation Site Analysis (ANSA), where discrete fluorescent amplification sites form and the number of sites correlates with input target concentration [32]. ANSA does not require partitioning into droplets or microwells and can be performed in either paper membranes or a simple microfluidic chip and can be imaged using a smartphone [32,33]. The assay readout is an image containing spatially localized, fluorescent amplification nucleation sites, and quantification requires detecting and counting individual sites. At higher target concentrations, these sites can grow and merge, causing standard counting methodologies to result in undercounts thereby narrowing the usable dynamic range. We hypothesize that the full image sequence recorded over 15 minutes contains additional quantitative information beyond site counts alone. The spatiotemporal evolution of site number, fluorescence intensity, and growth over time are not captured by conventional analyses that rely on explicit feature extraction such as intensity limits, threshold times, or discrete site counts. Convolutional neural networks (CNNs) provide a natural framework for learning hierarchical spatial features directly from imaging data and, when applied to multi-frame inputs, can capture spatiotemporal patterns relevant to classification. CNN-based models have shown strong performance across a wide range of medical image analysis tasks, including fundus imaging, computed tomography, radiology, and digital pathology [34–37].

In this paper, we develop a residual neural network (ResNet-18) [38] workflow to analyze time-resolved, multi-frame fluorescence images of RPA reactions within a microfluidic chip to semi-quantitatively classify HIV-1 DNA concentrations across five orders of magnitude and into four clinically relevant concentration categories. The models are trained on images acquired at fixed time points during each reaction enabling the model to learn both the spatial patterns and temporal evolution of fluorescent nucleation site formation. This approach avoids explicit site counting and threshold-based feature extraction, which become unreliable when ANSA nucleation sites overlap at high DNA concentrations. More broadly, these results demonstrate how machine learning can expand the quantitative utility of simple image-based isothermal amplification assays for decentralized molecular diagnostics.

## Methods

### Overview of ANSA Method and Image Analysis using ResNet Model

We apply a ResNet-18 convolutional neural network to classify time-resolved fluorescence images of ANSA reactions into either clinically defined HIV-1 DNA concentration ranges or logarithmically spaced concentration ranges. Two semi-quantitative classification models were developed: a clinical model with four output classes and a logarithmic model with five output classes.

For the clinical model, classes were defined as: undetectable (<200 copies), low (200–1,000 copies), medium (1,000–10,000 copies), and high (>10,000 copies). These class boundaries were based on established HIV-1 viral load thresholds used in clinical care, including the CDC threshold of <200 copies/mL for viral suppression[39–41] and the World Health Organization treatment monitoring threshold of <1,000 copies/mL[42–44]. In this study, these clinical viral load thresholds were used to define corresponding categories of input HIV-1 DNA copies per reaction for model training and classification.

For the logarithmic model, classes were defined as: <100 copies, 100–1,000 copies, 1,000–10,000 copies, 10,000–100,000 copies, and >100,000 copies. This formulation was used to separate samples across the full assay range without relying on disease-specific clinical thresholds. A classification approach was selected rather than regression because the dataset consisted of a limited number of samples at discrete concentration groups with relatively sparse sampling between concentrations. Figure 1 illustrates the process for semi-quantitative DNA measurement where fluorescence images of RPA reactions at different time points are fed into the trained ResNet-18 model, which outputs the approximate DNA concentration classes.

**Figure 1.**
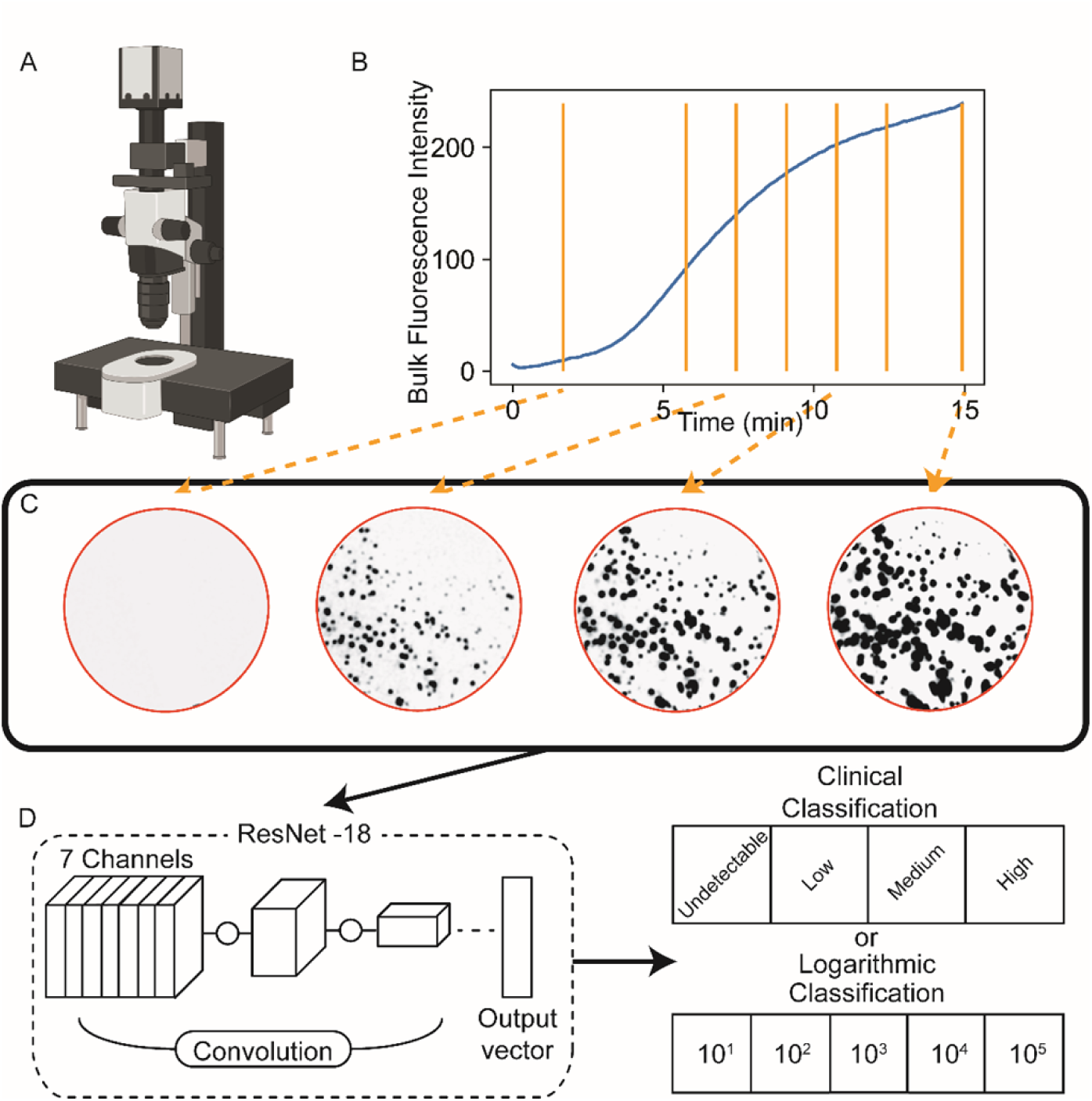
ANSA data analysis method using ResNet-18 models. (A) Time-resolved fluorescence microscopy images of ANSA reactions occurring in a microfluidic chip are acquired every 5 seconds over a 15-minute period. (B) Area averaged fluorescence intensity as a function of time showing a characteristic RPA sigmoidal amplification curve. Orange dashed lines denote seven representative frames selected as ResNet input channels. (C) Example microscopy frames after background subtraction and resizing (500 × 500 pixels). Four of the seven input frames are shown for clarity. Images are shown with inverted contrast for ease of visual interpretation, and the reaction area is outlined in red. (D) A ResNet-18 model was trained using labeled image sequences extracting spatiotemporal features to classify HIV-1 viral load across four clinical categories or spanning a 5-log DNA concentration range.

We perform digital isothermal nucleic acid amplifications in planar microfluidic chips, called Amplification Nucleation Site Analysis (ANSA). ANSA uses a viscous reaction buffer that results in discrete fluorescent amplification sites within a bulk solution. The ANSA reaction chemistry, loading of the microfluidic chip, and fluorescence imaging of the reactions have been described in our previous work [33]. Briefly, 100 μL RPA reactions containing target HIV-1 DNA were pipetted into the circular reaction chamber of a custom microfluidic chip, placed on a heating plate set to 39 °C, and imaged every 5 seconds over a 15-minute incubation using epifluorescence microscopy. The full sequence of images collected from a single reaction with a known concentration of DNA is referred to here as an image stack. A total of 273 separate ANSA reactions were conducted over an input DNA range of 20–10^6^ copies per reaction.

The mean pixel intensity of ANSA images as a function of time produces a sigmoidal fluorescence curve characteristic of RPA isothermal amplification dynamics, as shown in Fig. 1B. This profile mirrors conventional bulk-phase RPA reactions, with an initial lag phase, exponential signal increase, and subsequent plateau. Processing the entire image stack from each reaction is computationally expensive and ANSA reactions proceed over a 15-minute period thus the fluorescence image from one frame does not significantly differ substantially from the preceding and subsequent frames. To capture distinct stages along this trajectory, in a standardized way across all the experiments, seven key individual input images were selected as model input channels (Fig. 1B, vertical gold bars) to represent the major phases of the reaction.

The first input image was generated by averaging the first 20 frames to define a background baseline. The remaining six inputs were single frames taken at fixed time points: frame 50 (4:05 min; early growth at high concentrations), frame 70 (5:45 min; early growth at low concentrations), frame 90 (7:25 min; mid-reaction), frame 110 (9:05 min; early plateau), frame 130 (10:45 min; mid-plateau), frame 150 (12:25 min; late plateau), and frame 180 (15:00 min; reaction endpoint). In exploratory analyses, increasing the number of input channels to as many as 16 did not improve model performance (data not shown).

### Image Preprocessing and Dataset Preparation

The grayscale fluorescence TIFF image stacks were loaded using the scikit-image library (v0.24.0) [45]. Image acquisition and preprocessing were performed as previously described [33]. Briefly, each raw image contained two circular reaction areas, which were manually cropped into separate into individual square regions of interest, each centered on a single reaction area. A circular digital mask was then applied to each frame to isolate the reaction region and exclude peripheral areas of the chip. To reduce pixel-level noise, Gaussian blurring was smoothed using a 7 × 7 kernel. Global background subtraction was performed by computing the per-pixel temporal average of the first 25 frames and subtracting this background image from all frames, thereby setting the baseline background intensity approximately to zero and reducing static bright artifacts from the glass chip.

After preprocessing, each image stack was converted into a PyTorch[46] tensor, and resampled via bicubic interpolation to a uniform spatial resolution of 500 × 500 pixels across all frames. Resampling was required to correct minor dimensional inconsistencies introduced during manual cropping of raw images, ensuring consistent spatial input across all samples. No additional per-frame or per-stack intensity normalization, intensity clipping, or data augmentation was applied. Each image stack retained its DNA concentration label for training and was saved in a standardized PyTorch format. The dataset was then randomly shuffled at the individual reaction level and partitioned into training (60%), validation (20%), and test (20%) subsets as shown in Table 1 and Table 2. The test subset was not evaluated until hyperparameter selection was complete and the final model had been trained.

**Table 1.** Clinical utility model (Train/Validation/Test) split.

|  | Undetectable | Low | Medium | High |
| --- | --- | --- | --- | --- |
| Train | 44 | 29 | 26 | 60 |
| Validation | 14 | 13 | 13 | 18 |
| Test | 13 | 14 | 14 | 15 |

**Table 2.**
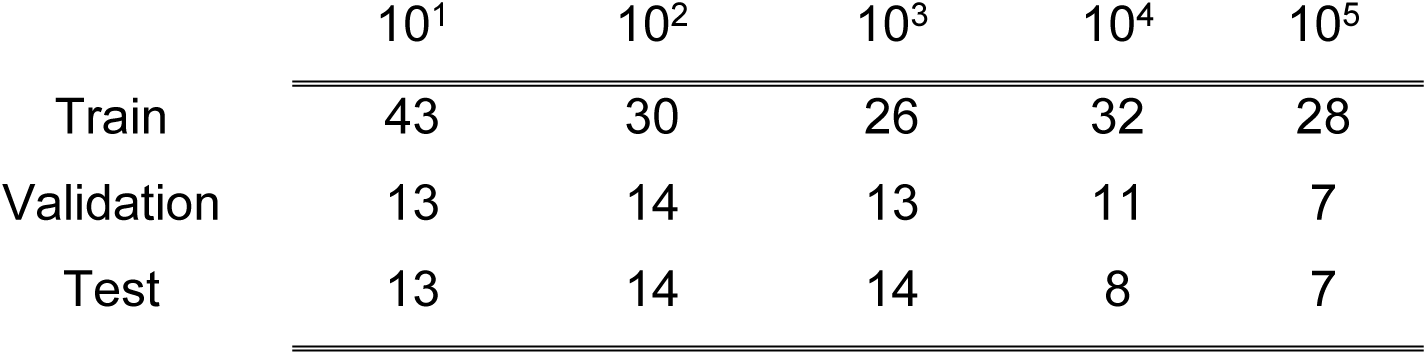
Logarithmic scale model (Train/Validation/Test) split.

### Model Architecture and Training

The selected model was chosen as a pragmatic approach for incorporating temporal image information into a standard 2D convolutional neural network. Temporally sampled fluorescent images from each ANSA reaction were provided to a modified ResNet-18 model as separate input channels by replacing the first convolutional layer to accept seven input channels, corresponding to the seven selected input images, rather than the default three channels used for standard RGB images used in the original ResNet. The fully connected classification head was replaced with a sequential module consisting of a dropout layer followed by a linear layer that output four DNA concentration classes in the clinical model or five DNA concentration classes for the logarithmic model. Training was performed using an Nvidia RTX 3060ti GPU with CUDA acceleration. Data were loaded using custom Dataset and DataLoader classes implemented in PyTorch v2.6.0+cu126 with a batch size of 32. To address class imbalance, a weighted random sampler was implemented based on inverse class frequency. The training set was used to update model weights during each training epoch, whereas the validation set was used to monitor model performance during training and guide hyperparameter optimization. Each model was tuned independently, and the final hyperparameters were selected separately for each model based on validation-set performance. Performance metrics including training loss, validation loss, and classification accuracy were recorded at each epoch. The held-out test set was not used for hyperparameter tuning or model selection and was evaluated only after the final model and hyperparameters had been selected. Final model performance was assessed on the test set, and a confusion matrix was generated to visualize classification performance.

### Hyperparameter Optimization

Hyperparameter tuning was performed using OPTUNA (v4.2.0)[47] with Bayesian optimization. The search space included model depth (ResNet-18 or ResNet-34 or ResNet-50), learning rate, weight decay, dropout rate, scheduler gamma value (the rate of exponential learning rate decay), and optimizer algorithm (Adam, SGD, ASGD, LBFGS). The optimization objective was to maximize the validation accuracy across 25 epochs, using a median pruning strategy to terminate underperforming trials early.

The model training initially used random weight initialization, which introduced substantial variability in validation performance across identical hyperparameter sets. An early version of the OPTUNA optimization loop addressed this by retraining each sampled hyperparameter set multiple times, retaining only the best-performing trial for each hyperparameter set before sampling new hyperparameters. This approach proved computationally inefficient and significantly slowed convergence, but improved robustness to random initialization effects. The final optimization strategy adopted a pruning and resampling approach. Trials were evaluated at intermediate epochs, and poorly performing configurations were terminated early. This strategy converged on similar optimal hyperparameters to those identified in the original retrain-and-select approach but did so more efficiently, although it required a greater total number of trials. For each model (clinical utility and logarithmic), 200 OPTUNA trials were conducted to identify the final hyperparameter set used for model training.

## Results

### ANSA reaction dynamics

We used real-time fluorescence imaging of ANSA reactions within a microfluidic chip to qualitatively observe how the fluorescence patterns evolved over time and varied with respect to input DNA concentration. Representative image sequences showed that reactions with higher input DNA produced more fluorescent puncta and reached higher area-averaged fluorescence intensities than lower concentration reactions (Figure 2). Low copy reactions showed delayed fluorescence onset and relatively few distinct nucleation sites. As input concentration increased, fluorescence appeared earlier and the number of visible sites increased, followed by progressive site growth and merging over time. At the highest concentrations, overlapping nucleation sites produced broad contiguous fluorescence rather than discrete puncta making site identification unreliable. Figure 2 shows these concentration-dependent differences in the spatial and temporal progression of amplification. We observed that DNA quantification with ANSA is limited when using endpoint bulk fluorescence, time-to-threshold, or site counting alone (see Figure S1) [33] We hypothesized that deep learning may extract spatiotemporal features from ANSA reactions that correlate with DNA concentration, and we implemented a ResNet-based analysis approach on real-time ANSA image sequences to provide robust semi-quantitative classification.

**Figure 2.**
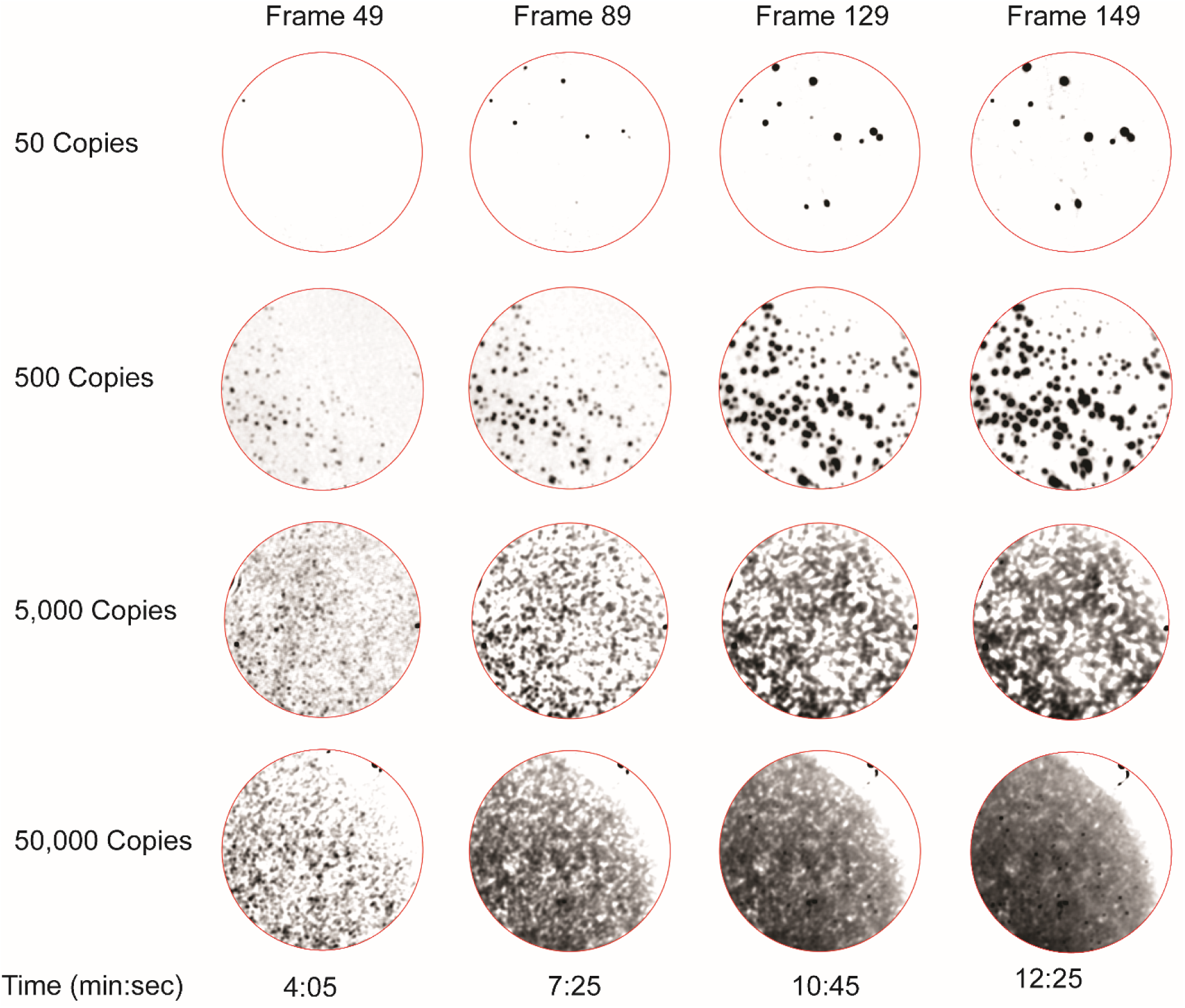
Experimental fluorescence images of ANSA reactions for 50, 500, 5,000, and 50,000 copies per reaction for the times listed (horizontal axis). Images are displayed with inverted contrast (ANSA sites are dark) for ease of interpretation, and the reaction area is outlined in red. As input DNA concentration increases, fluorescence appears earlier in the reaction, the number of visible nucleation sites increases, and site overlap becomes more pronounced over time. At the highest concentrations, overlapping sites produce broad contiguous fluorescence rather than discrete puncta, limiting reliable individual site counting.

### Model optimization and selected architecture

We perform hyperparameter optimization separately for the clinical and logarithmic classification tasks to identify model configurations that maximize ResNet classification performance. The clinical and logarithmic models were both trained for 25 epochs, but the optimal hyperparameters differed between tasks as shown in Table 3, indicating that the two classification problems imposed different learning demands. The clinical utility model converged with Stochastic Gradient Descent (SGD), a relatively higher learning rate, and moderate dropout, whereas the logarithmic model performed best with Adaptive Moment Estimation (Adam), a lower learning rate, and higher dropout. For both tasks, ResNet-18 performed comparably to deeper candidate architectures, including ResNet-34 and ResNet-50, and was therefore selected as the final model architecture.

**Table 3.** Summary of hyperparameters.

| Hyperparameter | Clinical Utility Model | Logarithmic Model |
| --- | --- | --- |
| Model Depth | 18 | 18 |
| Learning Rate | $3.88 \times 10^{-3}$ | $1.07 \times 10^{-4}$ |
| Weight Decay | $1.93 \times 10^{-4}$ | $2.37 \times 10^{-6}$ |
| Dropout Rate | 0.243 | 0.496 |
| Gamma Rate | 0.939 | 0.940 |
| Optimizer | SGD | Adam |

### Performance of the clinical utility model

Figure 3 shows the accuracy of the clinical utility ResNet classification model using a confusion matrix. The clinical utility model classified samples into four clinically relevant categories and achieved an overall test accuracy of 94.6%. The sensitivity was 100% in the undetectable, low, and high categories and lower in the medium category (78.6%). All errors in the medium category were restricted to adjacent classes, rather than representing large misclassifications across the concentration range. These results indicate that the model can accurately distinguish clinically relevant viral load categories while preserving appropriate category ordering in the few samples that were misclassified.

**Figure 3.**
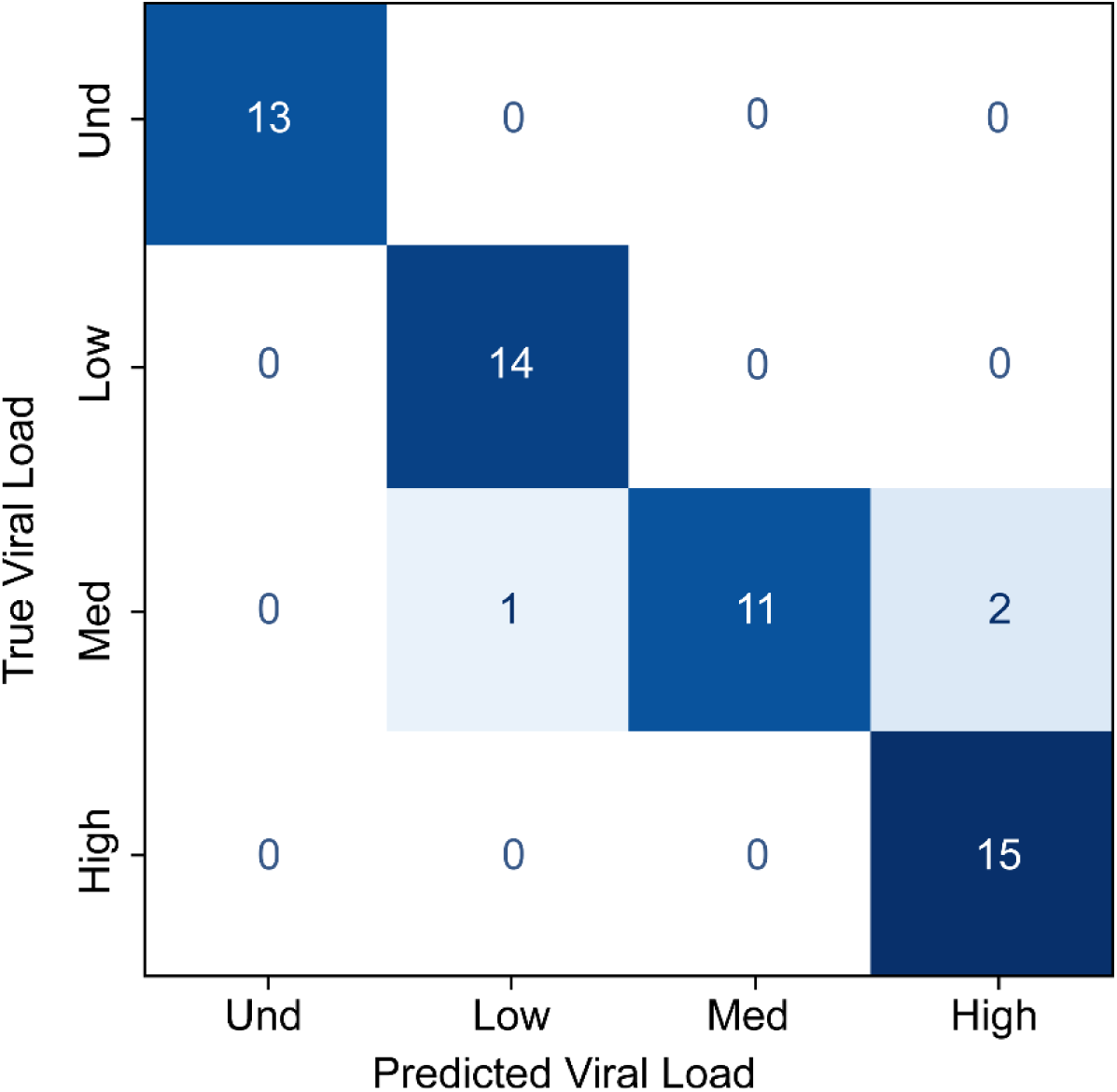
Clinical utility confusion matrix generated from the test dataset. The matrix displays predictions across four categories, with color intensity reflecting classification frequency. Diagonal entries represent correct predictions.

### Performance of the logarithmic model

Figure 4A shows the accuracy of the logarithmic ResNet classification model using a confusion matrix. The logarithmic model classified samples across five orders-of-magnitude bins and achieved an overall test accuracy of 92.7%. The sensitivity was 100% for 10⁴ and 10⁵ input copy number classes and remained high for the 10¹, 10², 10³ copy classes, 92.3%, 85.7%, and 85.7%, respectively. Figure 4B shows the true input copy numbers for each test sample grouped by predicted class, with misclassified sample explicitly marked. The logarithmic model misclassifications were also limited to adjacent bins, suggesting overlap between neighboring classes rather than systematic model error. These results demonstrate that ResNet-based analysis can support semi-quantitative nucleic acid classification across a broad dynamic range, including concentrations at which conventional ANSA site counting becomes unreliable.

**Figure 4.1.**
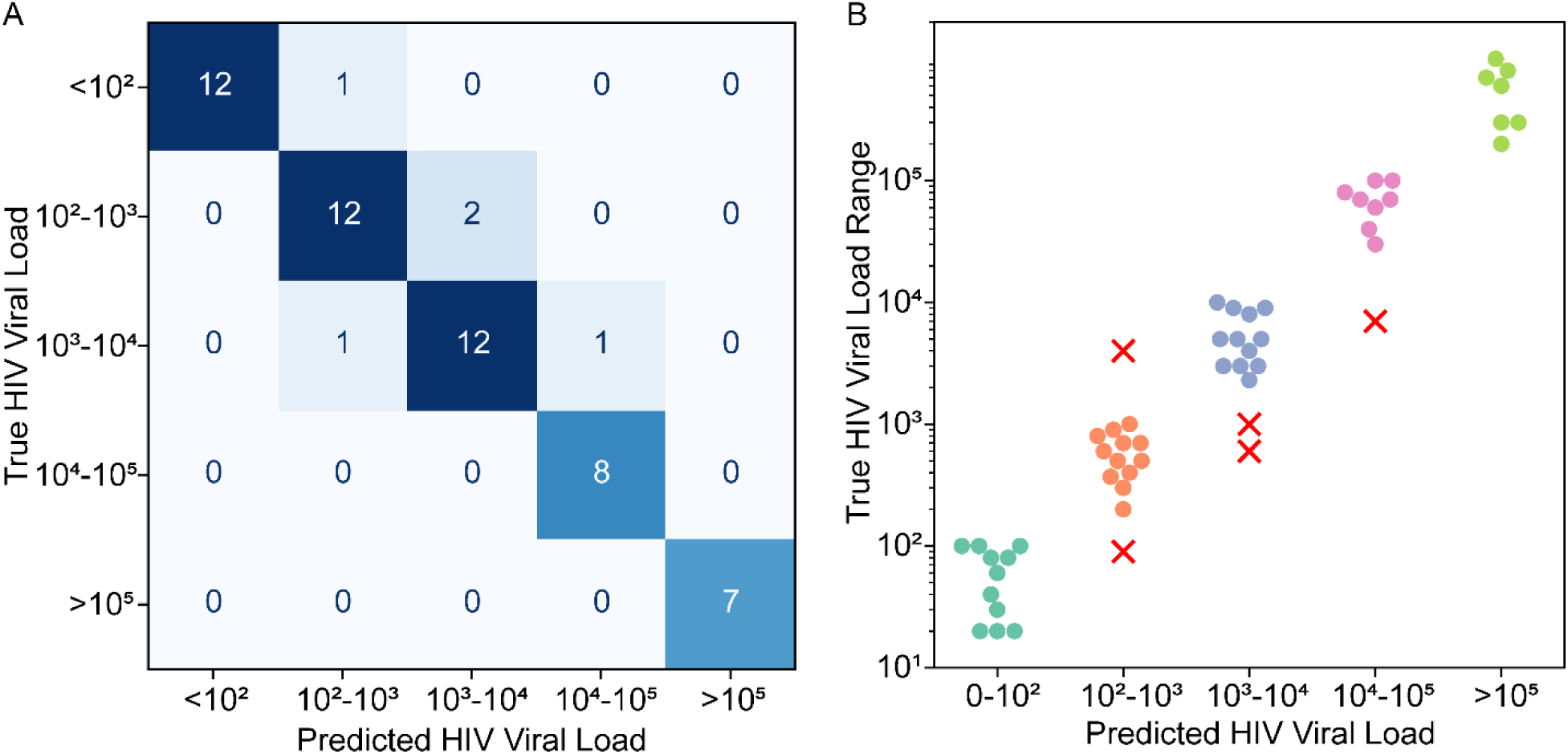
Classification performance using the logarithmic-scale model. (A) Confusion matrix showing classification performance across logarithmic-scale categories. Each cell represents the number of samples assigned to each predicted category. Diagonal entries indicate correct classifications, with darker shades representing higher counts. (B) True viral loads for each assay in the test dataset grouped by the predicted diagnostic category. Incorrect predictions are marked with a red X. Viral load is noted in total number of DNA per ANSA reaction.

## Discussion

Both clinical and logarithmic ResNet models accurately classify ANSA images using spatiotemporal image information. Clinically, nucleic acid measurements are often interpreted within broad categorical or log-scale ranges rather than as exact continuous values, despite the substantial concentration span within each decade. Consistent with that practice, our approach was designed as a semi-quantitative classifier. Rather than relying on direct puncta counting, the ML model uses the spatiotemporal evolution of fluorescence to assign reactions to concentration bins. This is especially advantageous for measuring high viral loads, where nucleation site overlap limits reliable direct counting, yet the ML model still preserved strong classification performance across nearly five orders of magnitude.

Although HIV clinical management guidelines are typically defined in units of viral load (copies per mL of plasma), the ANSA assay inputs in this study were defined as HIV-1 DNA copies per reaction, which is the standard analytical unit used during assay development and characterization. The clinical classification categories used here therefore represent input copy number ranges chosen to approximate established viral load thresholds rather than direct clinical measurements of viral loads from plasma samples. Future work toward HIV viral load monitoring will require a sample preparation step for extracting viral RNA plasma and recalibration using clinical specimens whose viral loads have been quantified using a gold standard clinical assay.

Most classification errors occurred at intermediate concentrations, particularly between 10^2^ and 10^3^ copies, where adjacent classes exhibited similar fluorescence progression and image morphology. These results show that these classes represent the most difficult decision boundary because their reactions share partially overlapping temporal and spatial features, including similar brightness evolution and less distinct separation in threshold timing. These errors reflect the similar fluorescence progression and image morphology of neighboring concentration groups rather than imaging artifacts or gross model instability.

ResNet-18 was selected as a relatively lightweight architecture that performed comparably to deeper candidate models during hyperparameter optimization, while maintaining lower computational complexity. In this implementation, temporally sampled fluorescence images were treated as separate input channels within a modified 2D CNN framework rather than using architectures explicitly designed for temporal data, such as 3D CNNs or recurrent neural networks. Although this approach does not directly encode temporal ordering, it was sufficient to capture joint spatiotemporal fluorescence patterns across reactions and enabled strong classification performance on the current dataset. The strong classification performance observed in both the clinical and logarithmic models suggests that these multi-timepoint fluorescence patterns contain sufficient information for robust semi-quantitative classification across a broad concentration range. Future work could evaluate dedicated spatiotemporal architectures to determine whether explicit temporal modeling further improves classification accuracy.

This study presents a machine learning approach for semi-quantitative classification of rapid isothermal nucleic acid amplification reactions. We developed a ResNet-18 image classification algorithm to quantify HIV-1 nucleic acids using real-time fluorescence imagery from diffusion-limited RPA reactions in a simple microfluidic chip. Using time-resolved images and a ResNet-18 model, we distinguished input DNA concentrations across up to five orders of magnitude.

This image-based classification approach may be useful for point-of-care molecular testing because it enables rapid, automated, and semi-quantitative interpretation of ANSA reactions within a low-cost chip. The logarithmic model may be especially useful for assays or pathogens that lack established clinical thresholds, whereas the clinical utility model provides output categories that are directly interpretable in an HIV monitoring context. At the same time, the current study remains a semi-quantitative classification approach rather than a continuous estimator, and ambiguity at intermediate concentrations remains an important limitation. Future work should evaluate larger datasets, additional targets, and deployment on compact or smartphone-based imaging platforms to determine how well this strategy generalizes across assays and use settings.

## Acknowledgments

This work was supported by the University of Washington eScience Institute’s Data Science & AI Accelerator Program, whose partnership with UW data scientists and domain researchers provided essential expertise, guidance, and infrastructure for our data-intensive experiments.

